# Rational Generation of Monoclonal Antibodies and Intrabodies Selective for Pathogenic TDP-43

**DOI:** 10.1101/2025.06.10.658846

**Authors:** Beibei Zhao, Sarah Louadi, Juliane A. Coutts, Ebrima Gibbs, Megan Y. Huang, Ryan J. Marina, Stefan Aigner, Parvez Alam, Mari L. DeMarco, Ian R. Mackenzie, Anke A. Djikstra, Byron Caughey, Gene W. Yeo, Johanne M. Kaplan, Neil R. Cashman

## Abstract

TAR DNA-binding protein 43 (TDP-43), encoded by the *TARDBP* gene, is a ribonucleoprotein associated with the pathogenesis of amyotrophic lateral sclerosis (ALS), frontotemporal dementia (FTD), and Alzheimer’s disease (AD). Under physiological conditions, TDP-43 is predominantly localized in the nucleus, where it participates in a variety of cellular functions related to RNA splicing, transport, and stability, as well as miRNA biogenesis. In disease, it is disproportionately mislocalized to the cytoplasm where it forms aggregates, which contribute to neurotoxicity and prion-like cell-to-cell propagation of pathogenic TDP-43. Targeting of misfolded aggregates of TDP-43 represents an attractive therapeutic strategy. However, development of effective immunotherapeutic agents remains a challenge, as they require stringent selectivity for misfolded TDP-43 in order to maintain the essential functions of physiologically native TDP-43. To address this issue, monoclonal antibodies (mAbs) and intrabodies were generated against an epitope in the N-terminal domain of TDP-43 that is only exposed when the protein is misfolded, but not in its properly folded form. We show that mouse and rabbit mAbs against this epitope displayed high binding affinities by surface plasmon resonance analysis and selectively reacted with pathological TDP-43 in post-mortem tissues from ALS, FTD, and AD patients. In a cell line system, human embryonic kidney (HEK) 293T cells, mAbs and corresponding intrabodies specifically reacted with cytoplasmic aggregates of transfected misfolded TDP-43 lacking the nuclear localization signal, TDP-43^ΔNLS^. Functionally, mAbs inhibited cell-to-cell transmission of misfolded TDP-43 and the seeding activity of misfolded TDP-43 from FTLD brain homogenates by a novel RT-QuIC assay. Intrabodies promoted the degradation of intracellular aggregates of TDP-43 in HEK293T cells and in induced pluripotent stem cell-derived motor neurons (iPSC-MNs) from ALS patients. The results provide proof-of-concept evidence that supports selective targeting of misfolded toxic aggregates of TDP-43 as a potentially safe and effective avenue to treat neurodegenerative diseases associated with TDP-43 proteinopathy.

## Introduction

Transactive response DNA binding protein 43 (TDP-43) is a heterogeneous nuclear ribonucleoprotein (hnRNP) encoded by the *TARDBP* gene ubiquitously expressed in most cells and tissues. Structurally, it consists of an N-terminal domain (NTD), a nuclear localization signal (NLS), two RNA-recognition motifs (RRMs), two intrinsically disordered domains (IDRs), and a low complexity domain (LCD; glycine-rich C-terminal) (1).

TDP-43 plays a crucial role in various cellular events, such as transcriptional regulation, splicing, mRNA transport and stability, stress response, and neuronal-specific protein expression (2). Under physiological conditions, TDP-43 shuttles between the nucleus and cytoplasm, with ∼90% being localized to the nucleus at any given time (3, 4). However, under disease conditions, such as amyotrophic lateral sclerosis (ALS), frontotemporal dementia (FTD), and Alzheimer’s disease (AD), TDP-43 is frequently depleted from the nucleus and mislocalized to the cytoplasm where it forms aggregates in affected neurons and glial cells. These pathogenic TDP-43 aggregates are believed to be a primary driver of disease through gain-of-function (GoF) and/or loss-of-function (LoF) (4), which contribute to direct neurotoxicity within the cell, and prion-like propagation from cell to cell and from region to region in the central nervous system (CNS) (4–12).

ALS is the most common motor neuron disease characterized by progressive degeneration of motor neurons in the spinal cord and motor cortex of the brain that control limb movement, swallowing, speech, and breathing. With disease progression, ALS ultimately leads to motor neuron death, resulting in muscle paralysis and respiratory failure. Approximately 10% of ALS is familial (fALS) associated with various genetic mutations. The remaining ∼90% is sporadic (sALS). FTD is characterized by neuronal degeneration in the frontal and temporal lobes of the brain. The most common symptoms include behavioral changes, language disorders, and in some cases motor system degeneration as seen in ALS. ALS and FTD are considered to belong to a common disease spectrum (ALS–FTD) because of overlaps in genetics, clinical symptoms, and pathologies (13). Accumulation of TDP-43 inclusions in the cytoplasm is a major pathological hallmark shared by both diseases and is observed in ∼97% of all ALS and ∼50% of FTD (14, 15). TDP-43 proteinopathy has also been observed in up to 57% of Alzheimer’s disease (AD) (16), a progressive neurodegenerative disorder characterized by cortical degeneration. Interestingly, when AD with mixed pathology of limbic-predominant age-related TDP-43-encephalopathy (LATE) is observed on post-mortem neuropathology, it is associated with more rapid dementia in life (17). Targeting of pathogenic TDP-43 represents a potentially effective therapeutic approach, which necessarily comes with a requirement for isoform specificity of targeting misfolded TDP-43 while sparing normally folded healthy TDP-43 to perform its numerous physiological functions in a wide range of cellular events.

Immunotherapeutics, such as conventional antibodies and intracellular antibodies (intrabodies), have been explored in pre-clinical studies by a number of groups in an attempt to develop therapeutic strategies for the treatment of neurodegenerative diseases, including Huntington’s disease (HTT), Parkinson’s disease (PD), prion diseases, AD and ALS, and have shown promising results (18–23). Antibodies offer the potential to inhibit cell-to-cell transmission of pathogenic proteins associated with neurodegenerative diseases, thereby preventing the spread of neurodegeneration. Our previous studies have successfully demonstrated the ability of misfolding specific antibodies against defined epitopes exposed by protein misfolding to achieve efficacy and safety by specifically targeting the misfolded isoforms relevant to disease progression. Protective activity was observed with misfolding-specific passive and active immunotherapies including an amyloid-beta (Abeta) oligomer-specific antibody (24, 25), alpha-synuclein oligomer-specific antibodies (20), antibodies directed against pathogenically misfolded superoxide dismutase type 1 (SOD1) to inhibit *in vitro* cell-to-cell transmission of misfolded SOD1 (23, 26) associated with fALS-SOD1 cases, and wild-type SOD1 in sALS induced to misfold by physical contact with TDP-43 (22), as well as the use of misfolding-specific epitopes as immunogens in vaccines (27). Intrabodies, such as single-chain variable fragment (scFv) intrabodies, delivered *via* gene therapy vectors also have the potential to promote the clearance of pre-existing intracellular aggregates of pathogenic proteins (18).

Development of immunotherapeutics targeting pathogenic TDP-43 requires stringent selectivity of antibodies for misfolded TDP-43 to ensure safety. To address this challenge, in the present study, we developed monoclonal antibodies (mAbs) directed against a peptide epitope exposed by a loss of structure in TDP-43 N-terminal domain that selectively target misfolded/pathological TDP-43, while sparing normally folded healthy TDP-43 from immune recognition. mAbs and intrabodies against this epitope were screened for reactivity against pathological aggregates of TDP-43 in post-mortem tissues from ALS, FTD, and AD patients, and in HEK293T cells transfected with TDP-43^ΔNLS^. mAbs were also tested in a TDP-43 novel real-time quaking-induced conversion (RT-QuIC) assay, in which they showed inhibition of seeding propagation from FTLD-TDP brain homogenates. Activity of intrabodies was assessed in the HEK293T system, as well as in induced pluripotent stem cell-derived motor neurons (iPSC-MN) from ALS patients.

## Results

### Generation of mAbs

Tryptophan-68 (Trp68) in the NTD of TDP-43 is not available for interaction at the natively folded TDP-43 molecular surface, and is buried in the native structural core of the NTD (28). Partial unfolding *in silico* using Collective Coordinates (EpiSelect^TM^) (29) indicated that Trp68 is solvent-exposed when the TDP-43 protein structure is subjected to progressive loss of native atomic contacts, and our experimental research confirmed that Trp68 is exposed in misfolded TDP-43 (22). Polyclonal IgG from rabbits immunized with a seven amino acid peptide centered around Trp68 in the context of its local amino acid sequence (_65_DAGWGNL_71_) showed selective binding to misfolded, pathogenic TDP-43 and did not react with normally folded TDP-43 (22). In the current study, the same immunogen was used to generate monoclonal antibodies (mAbs) in mice and rabbits.

### TDP-43 mAbs display high binding affinity to a misfolding specific TDP-43 epitope

In order to selectively target pathogenic TDP-43 without interfering with the function of normal TDP-43, mouse and rabbit mAbs were raised against an N-terminal epitope that we previously determined to be restricted to misfolded TDP-43 (22). SPR analysis was performed to measure the binding affinity of candidate mAbs to this epitope. As shown in **Figure 1** and **Table 1**, the mAbs displayed high affinity for the epitope in the sub-nanomolar range, with rabbit mAbs displaying overall greater affinity than mouse mAbs.

**Figure 1.**
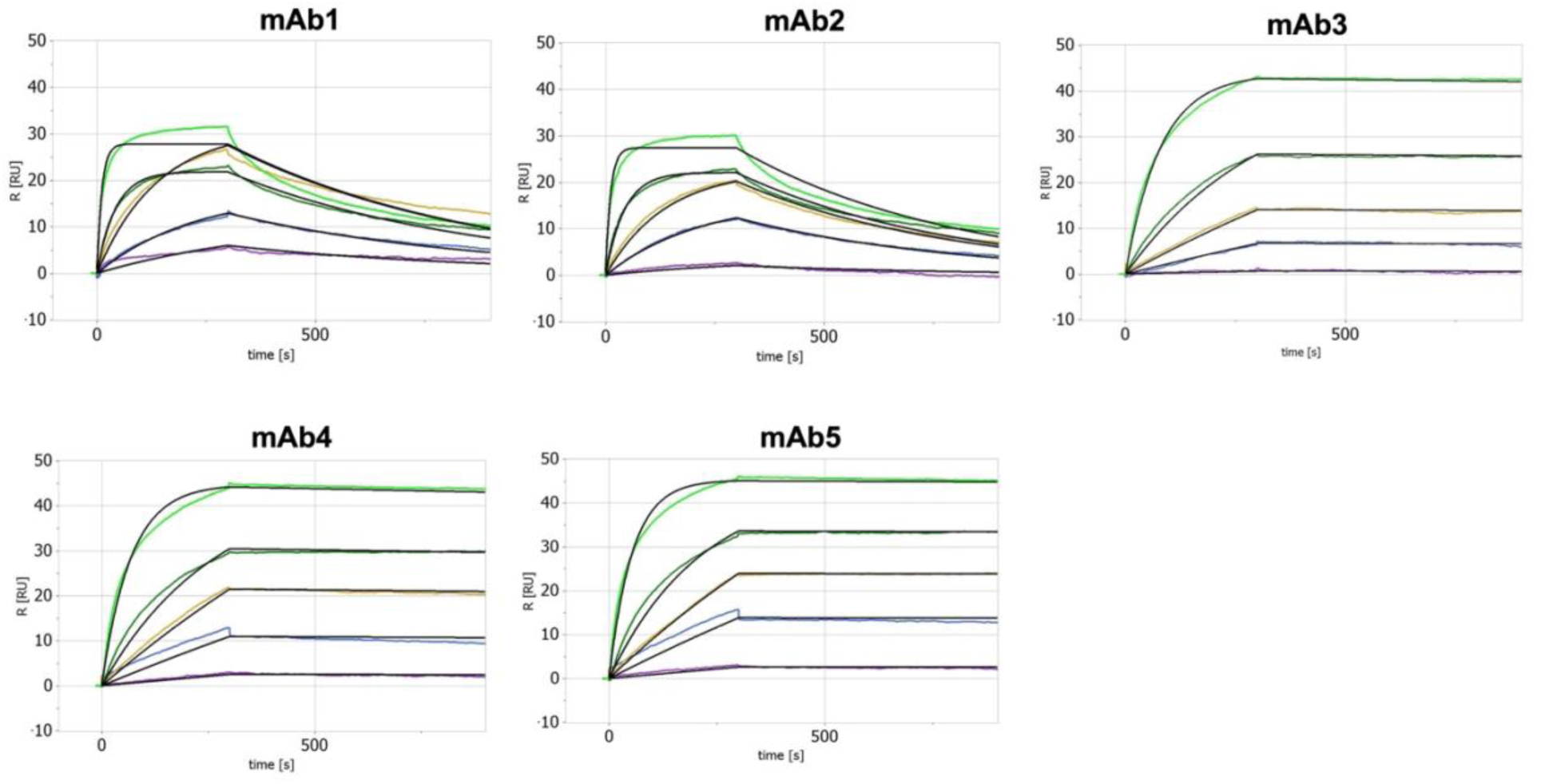
TDP-43 mAbs display high binding affinity to a misfolding specific TDP-43 epitope. SPR analysis shows high binding affinity of TDF-43 mAbs to the “_65_DAGWGNL_71_” epitope.

**Table 1.** Kinetic analysis shows TDP-43 mAbs display high binding affinity to the misfolded TDP-43 epitope in the sub-nanomolar range. mAb1 and mAb2 are mouse mAbs. mAb3 – mAb5 are rabbit mAbs. mAb1 and mAb5 were selected for further characterization.

| mAb | $k_{ON}$<br>($M^{-1}s^{-1}$) | $k_{OFF}$<br>($s^{-1}$) | $K_D$<br>(M) |
| --- | --- | --- | --- |
| mAb1 | 2.83E+06 | 1.76E-03 | 6.22E-10 |
| mAb2 | 2.77E+06 | 2.00E-03 | 7.22E-10 |
| mAb3 | 4.03E+05 | 2.50E-05 | 6.20E-11 |
| mAb4 | 4.70E+05 | 4.32E-05 | 9.19E-11 |
| mAb5 | 5.99E+05 | 9.04E-06 | 1.51E-11 |

### TDP-43 mAbs specifically recognize cytoplasmic aggregates of mutant TDP-43 in HEK293T cells

Two representative mAbs (one rabbit and one mouse) were selected for further analysis. To assess selectivity for pathogenic TDP-43, we next characterized the immunoreactivity of mAbs with misfolded TDP-43 in human embryonic kidney (HEK) 293T cells transiently transfected with an engineered HA-tagged nuclear localization signal (NLS)-deficient mutant of TDP-43, TDP-43^ΔNLS^, which forms cytoplasmic aggregates of misfolded TDP-43 detectable by immunocytochemistry (ICC). As shown in **Figure 2**, both antibodies specifically recognized aggregates of TDP-43^ΔNLS^ in the cytoplasm, with no detectable reactivity against normal endogenous nuclear TDP-43 in transfected cells or neighbouring un-transfected (UT) cells, demonstrating the selectivity of the mAbs for misfolded/aggregated TDP-43.

**Figure 2.**
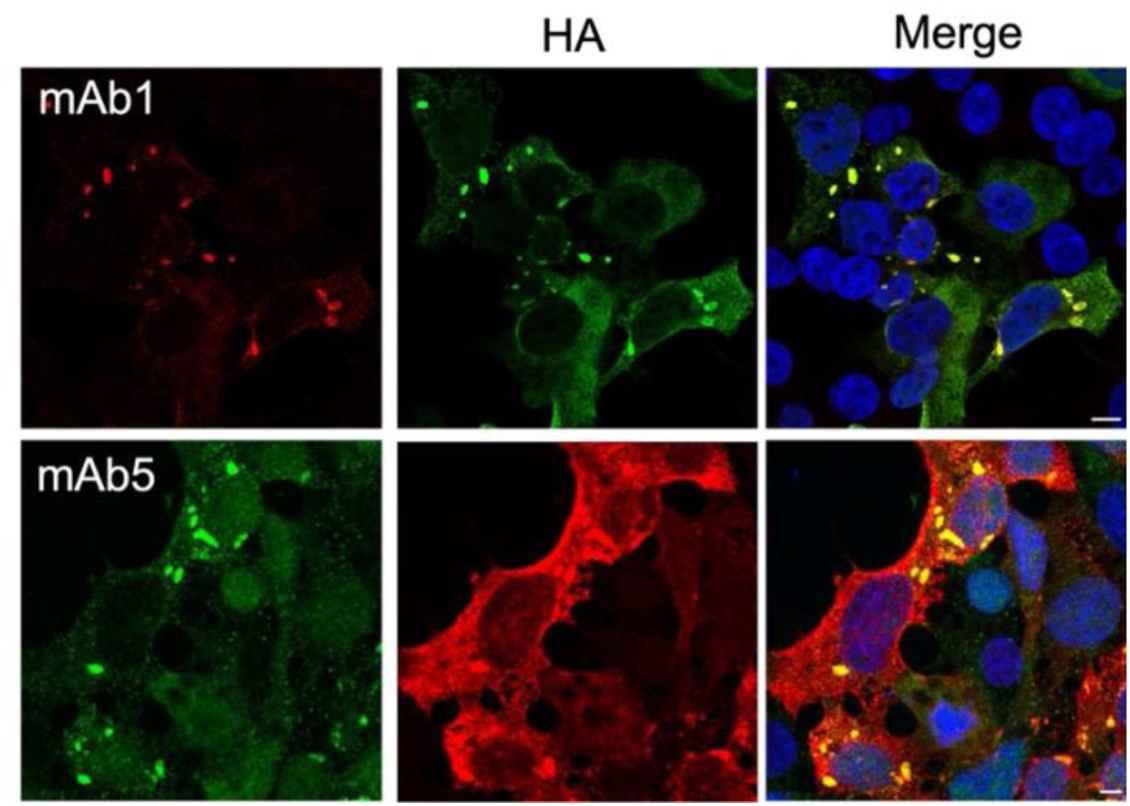
TDP-43 mAbs specifically recognize cytoplasmic aggregates of mutant TDP-43 in HEK293T cells. Immunocytochemical analysis demonstrates mAb1 and mAb5 interact specifically with cytoplasmic aggregates of TDP-43^ΔNLS^ (HA) in transfected HEK293T cells. Scale bars: 10 μm.

### TDP-43 mAbs do not recognize TDP-43 oligomers in physiological stress granules

Stress granules (SGs) are dynamic cytoplasmic membrane-less structures consisting of ribonucleoprotein (RNP) complexes formed in response to environmental stressors, such as oxidative stress, heat shock, nutrient deprivation, and viral infection (30). TDP-43, as an RNP protein, plays an important role in SG formation, which serves as a protective mechanism under stress conditions by storing untranslated mRNAs, temporarily arresting protein synthesis, and conserving energy. It has been shown that normal TDP-43 is recruited to physiological SGs upon acute oxidative stress (31–34). In contrast, pathological TDP-43 aggregates, such as those associated with ALS mutations, are typically not present in physiological SGs (30, 35). It has also been shown in *in vivo* studies that mutant TDP-43 impairs SG assembly and function (36). To assess the ability of our TDP-43 misfolding-specific antibodies to distinguish between TDP-43 in physiological SGs and pathological aggregates, HEK293T cells were treated with 1 mM sodium arsenite for 60 min (acute oxidative stress), and co-stained with mAb1 or mAb5 and an established SG marker, G3BP1. As shown in **Figure 3**, SGs were formed upon sodium arsenite treatment and staining with a pan TDP-43 antibody displayed the expected recruitment of TDP-43 to SGs. Neither mAb1 nor mAb5 reacted with these SGs, suggesting that the NTD of wild-type TDP-43 molecules recruited to SGs in cells that do not possess ALS-associated mutations maintain their physiological fold needed for functional oligomerization.

**Figure 3.**
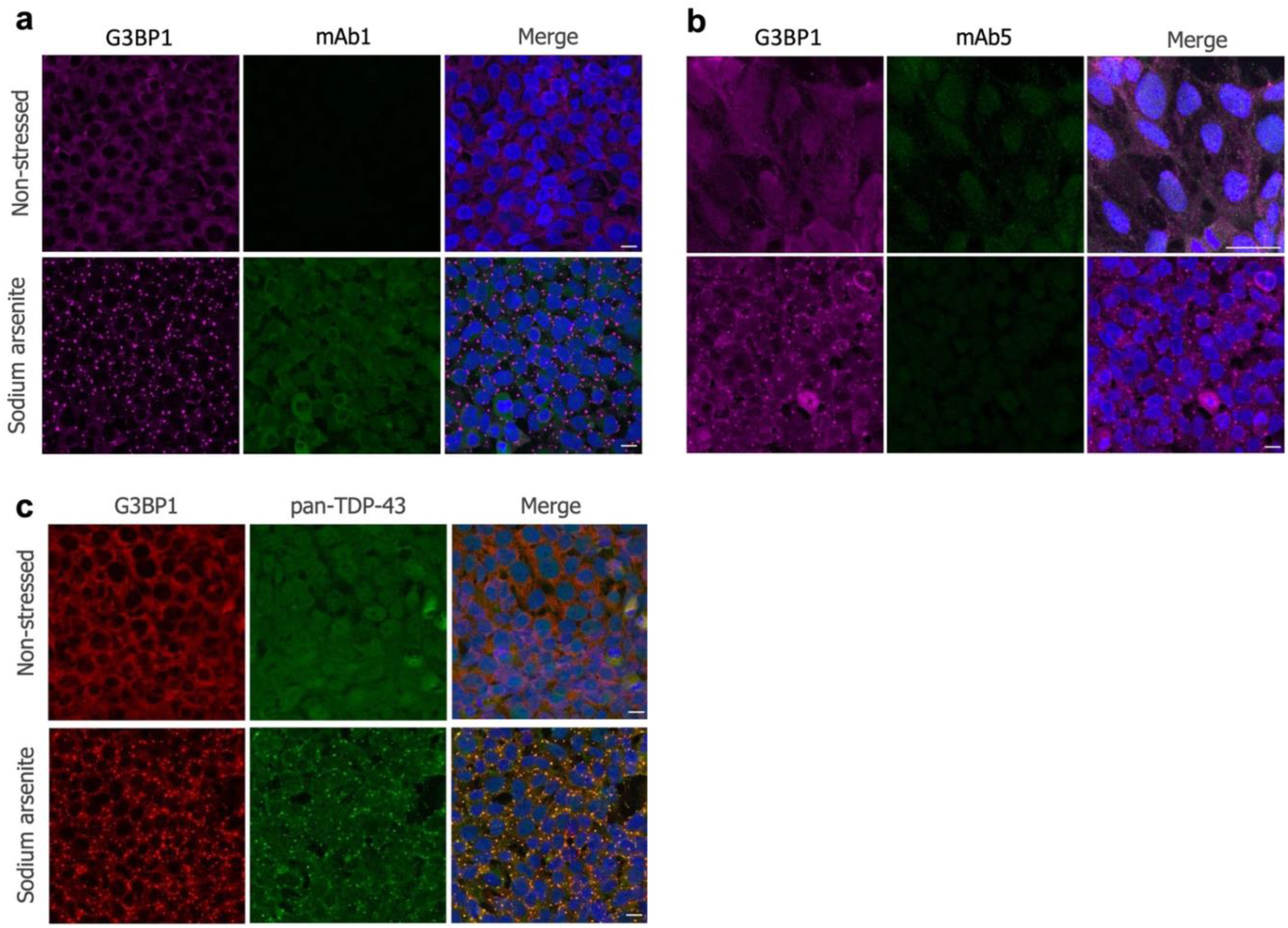
TDP-43 mAbs do not recognize oligomers of TDP-43 in physiological SGs. Immunocytochemical analysis demonstrates mAb1 and mAb5 did not recognize TDP-43 in physiological SGs (G3BP1) upon acute oxidative stress by sodium arsenite treatment (1 mM, 60 min) (**a, b**). A pan-reactive TDP-43 antibody confirms that TDP-43 is present in physiological SGs (**c**).

### TDP-43 mAbs detect pathological TDP-43 in ALS, FTD, and AD

Considering that TDP-43 pathology is present in 97% of ALS, 50% of FTD, and 57% of AD as mentioned above, we next characterized the immunoreactivity of the mAbs in post-mortem human tissues of ALS, FTD, and AD patients by immunohistochemistry (IHC). As shown in **Figure 4a**, cytoplasmic inclusions of TDP-43 were observed with mAb1 in both white and grey matter of an FTD-TDP type B case, similar to the IHC pattern of a commercial antibody against phosphorylated TDP-43 (pTDP-43), which is a hallmark of TDP-43 pathology in affected CNS regions of ALS, FTD, and AD patients (4, 15, 16, 37–39). Similarly, mAb5 specifically recognized thread-like and/or round inclusions of TDP-43 in the cytoplasm in ALS spinal cord, and FTD-TDP and AD brain tissues, with no immunoreactivity in healthy control samples (**Fig. 4b-d**). This is consistent with the staining pattern that we observed with a rabbit polyclonal antibody raised against the same epitope (22). By comparison, a commercial pan-TDP-43 antibody recognized both normal nuclear and cytoplasmic inclusions of TDP-43 (**Fig. 4b-d**). These results demonstrate the specificity of the mAbs for misfolded/pathological TDP-43.

**Figure 4.**
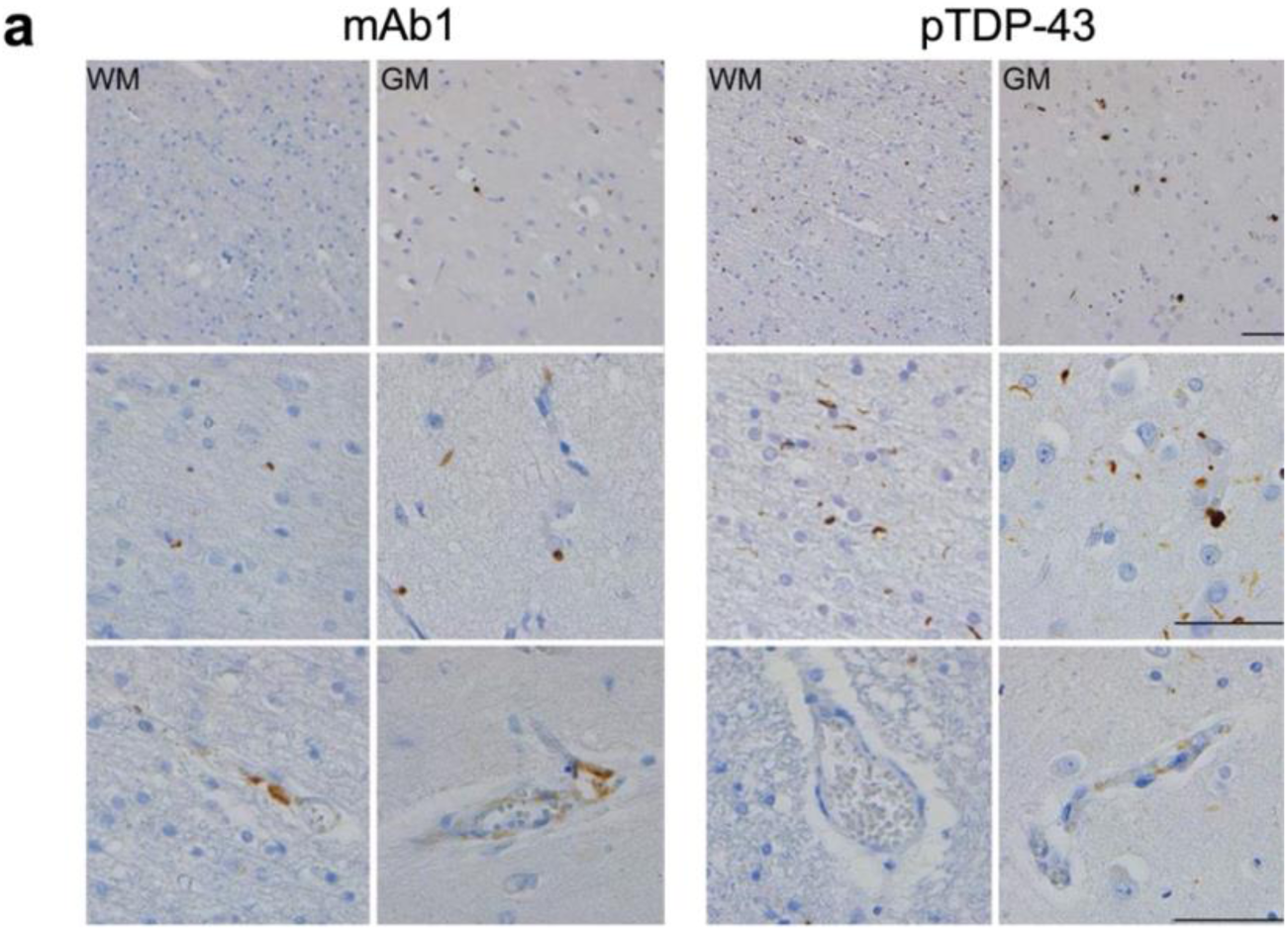
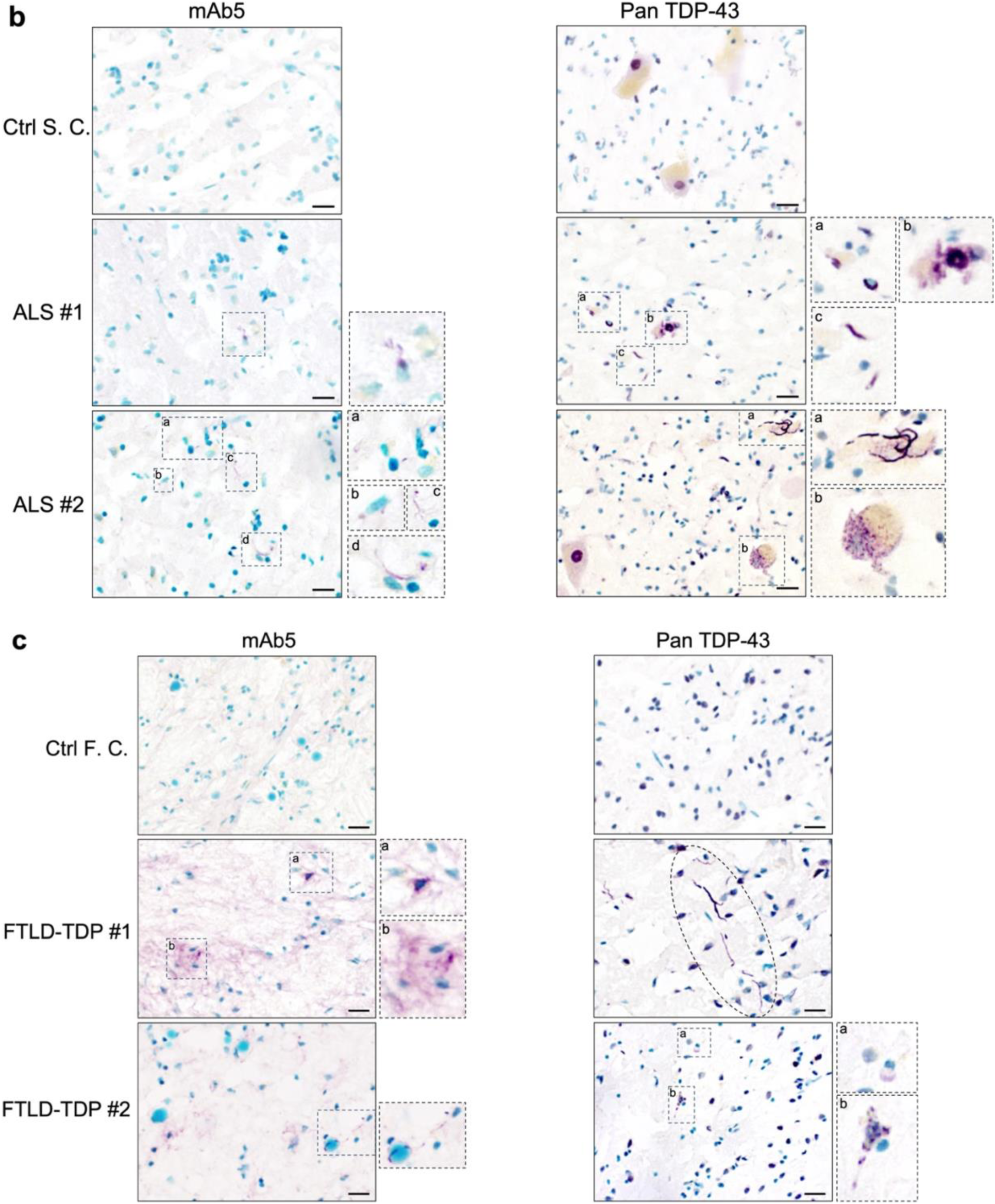
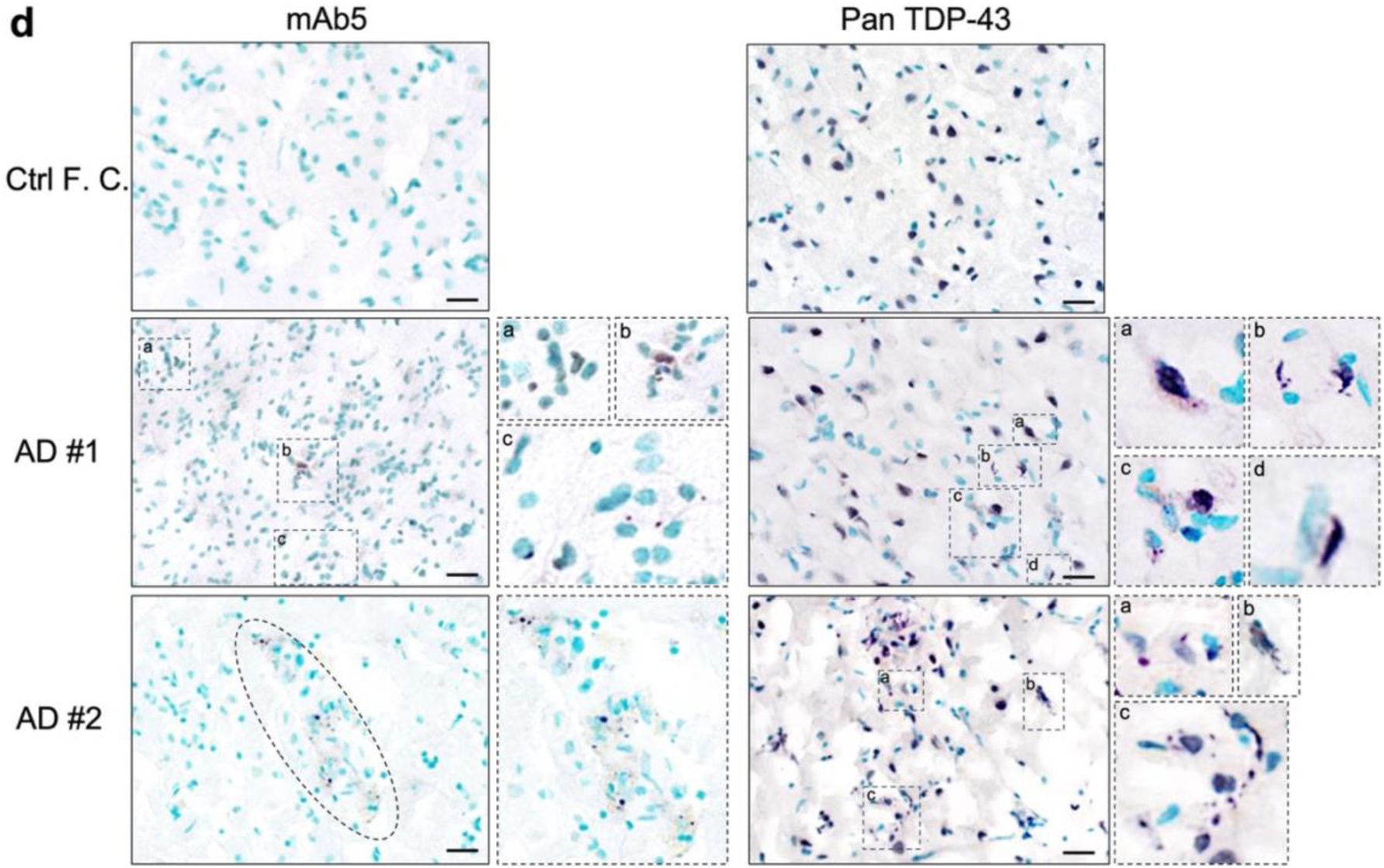
TDP-43 mAbs detect pathological TDP-43 in ALS, FTD, and AD. Immunohistochemical analysis demonstrates mAb1 detects thread-like and/or round inclusions of TDP-43 in the cytoplasm and deposits around blood vessels in the white matter (WM) and grey matter (GM) in an FTD-TDP type B case, resemblant to the staining pattern of phosphorylated TDP-43 (pTDP-43) (**a**). mAb5 specifically recognizes thread-like and/or round inclusions of TDP-43 in the cytoplasm in ALS spinal cords (SC, **b**), FTD-TDP and AD frontal cortex brain tissues (FC, **c&d**), with no immunoreactivity in healthy control (Ctrl) samples. A pan-TDP-43 antibody reacts with both normal nuclear TDP-43 in healthy controls and cytoplasmic inclusions of TDP-43 in ALS, FTD-TDP and AD (**b-d**). Scale bars: 50 μm

### TDP-43 mAb inhibits TDP-43^ΔNLS^ intercellular transmission in HEK293T cells

Mounting evidence has suggested that misfolded TDP-43 can be transmitted in the extracellular space from cell to cell and from region to region in the CNS in a prion-like fashion contributing to the spread of neurodegeneration (4, 5, 7–12). We therefore tested the potential of inhibiting cell-to-cell transmission with a mAb against misfolded TDP-43^ΔNLS^. We employed our previously published procedure (23), in which conditioned medium from donor cells transfected with TDP-43^ΔNLS^ was incubated with mAb1 or control IgG to allow for antibody binding to any misfolded TDP-43^ΔNLS^ aggregates released into the medium by the donor cells. The resulting medium was then added to naïve recipient cells for quantification of the ability of the antibody to inhibit TDP-43^ΔNLS^ transmission to recipient cells by western blotting (WB) analysis. As shown in **Figure 5**, mAb1 significantly reduced the amount of TDP-43^ΔNLS^ transmitted from donor to recipient cells compared to control IgG.

**Figure 5.**
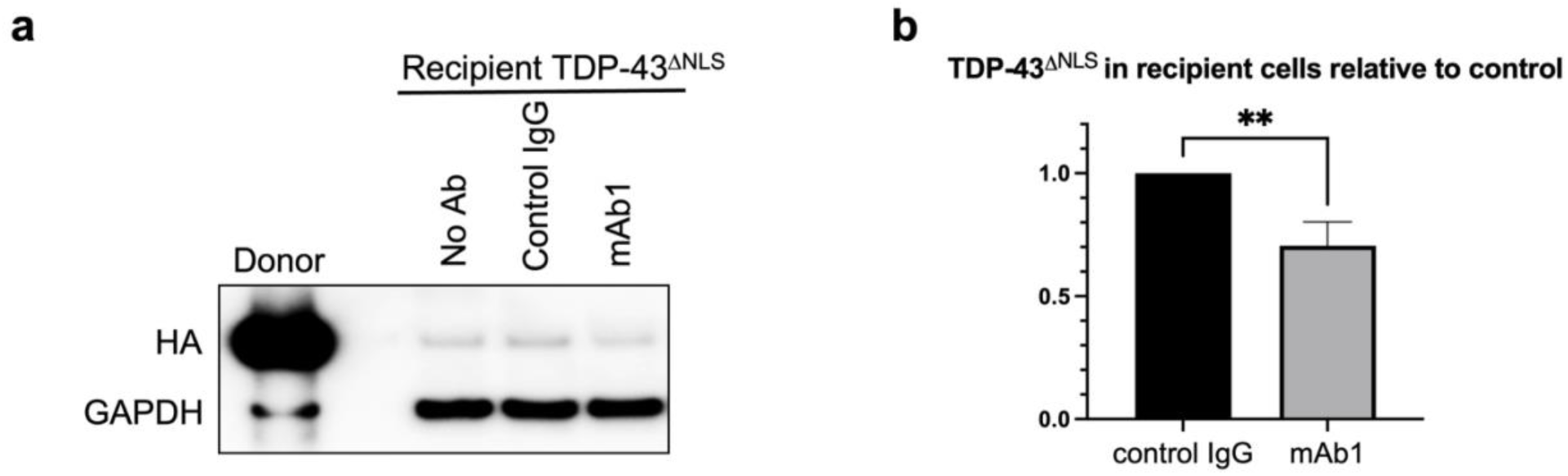
TDP-43 mAb inhibits TDP-43^ΔNLS^ intercellular transmission in HEK293T cells. Representative WB (**a**) and quantification (**b**) show mAb1 reduces transmission of TDP-43^ΔNLS^ (HA) from donor to naïve recipient cells compared to control IgG. GAPDH serves as the loading control. *Statistics*, Student’s *t*-test, unpaired two-tailed. \*\**p ≤* 0.01. n=3

### TDP-43 mAbs inhibit seeding of pathological TDP-43

We next tested whether TDP-43 mAbs could inhibit the prion-like seeding activity of misfolded TDP-43. To this end, we employed a TDP-43 assay using the well-established method Real-Time Quaking-Induced Conversion (RT-QuIC), a highly sensitive assay used to detect misfolded proteins associated with neurodegenerative diseases, including prion disease, Parkinson’s disease, ALS, and FTD (40–48). Our assay was developed independently of a previously reported TDP-43 RT-QuIC assay (40, 49). As shown in **Figure 6**, both mAb1 and mAb5 inhibited the seeding activity of misfolded TDP-43 derived from the brain homogenates from FTLD-TDP patients, with no reactivity with normal control samples, and no FTLD-TDP RT-QuIC inhibition with isotype control mAbs. Notably, the RT-QuIC seeding substrate was a fragment of the TDP-43 C-terminal low complexity domain (amino acids 263-414) lacking the NTD, suggesting that the seed neutralization of our mAbs was due to binding of pre-existing NTD-unfolded TDP-43 in the FTLD-TDP brain homogenates.

**Figure 6.**
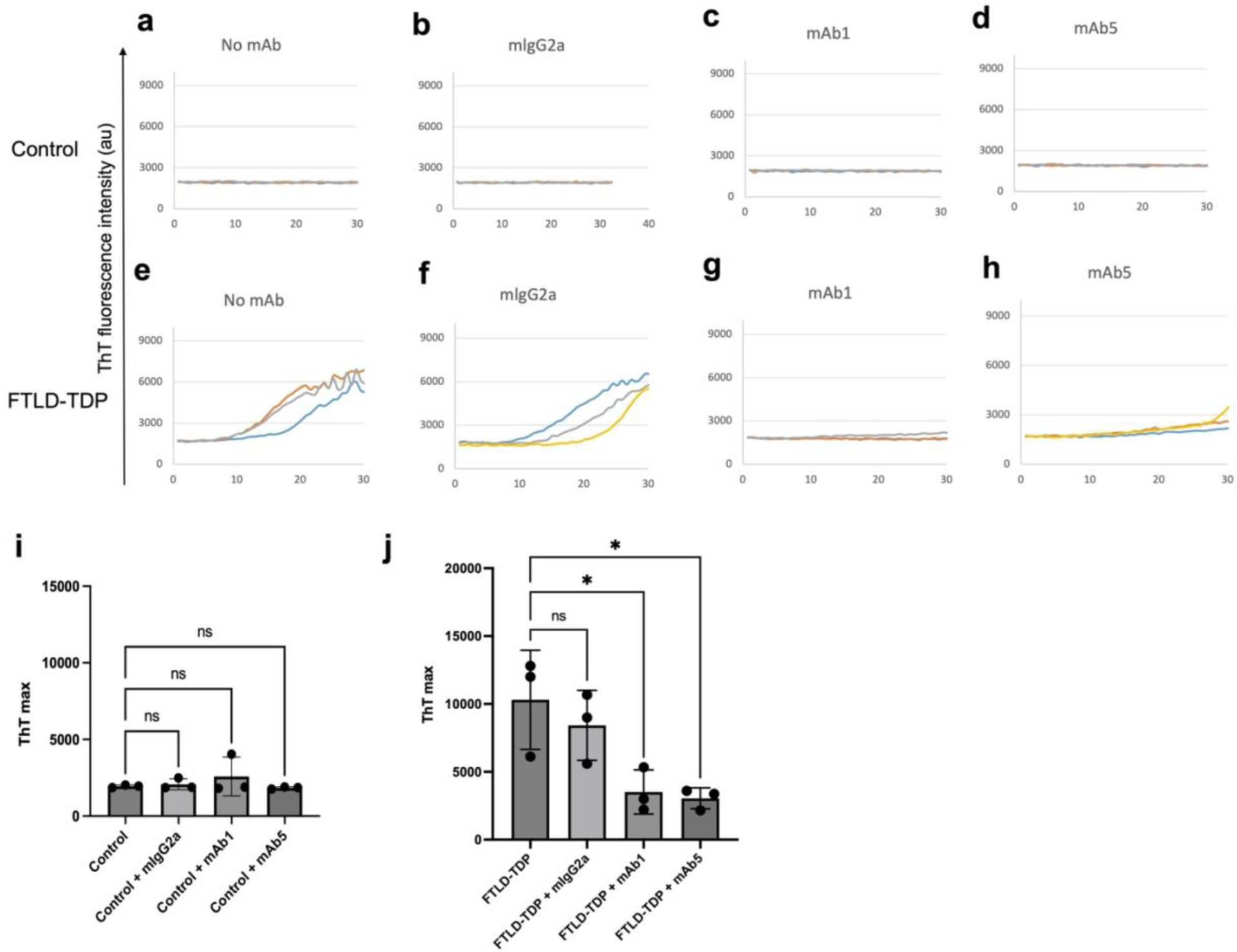
TDP-43 mAbs inhibit the seeding activity of FTLD-TDP brain homogenate in RT-QuIC assay. Representative TDP-43 RT-QuIC reactions in the absence **(a, e)** and presence of isotype control **(b, f),** mAb1 **(c, g)**, and mAb5 **(d, h)** with control **(a-d)** and FTLD-TDP brain samples **(e-h)**. Quantification of the ThT max of TDP-43 RT-QuIC reactions seeded with brain homogenates from control **(i)** and FTLD-TDP **(j)**. *Statistics*, one-way ANOVA. \**p* < 0.05, n=3.

### TDP-43 intrabodies recognize TDP-43^ΔNLS^ cytoplasmic aggregates and accelerate TDP-43***^ΔNLS^*** degradation

Intrabodies offer the advantage of targeting pathogenic proteins inside of the cell with the potential to clear pre-existing cytoplasmic aggregates of pathogenic TDP-43 and prevent further spread. To test this premise, we generated single-chain variable fragment (scFv) intrabody plasmid constructs encoding the variable binding regions of the heavy (VH) and light (VL) chains of mAb1 or mAb5 joined by a linker, with or without a lysosomal targeting signal as described in Materials and Methods (PMN A, PMN C, and PMN E).

First, to characterize the expression and intracellular binding profile of the intrabodies, ICC was performed in HEK293T cells co-transfected with TDP-43^ΔNLS^ and PMN A, PMN C, or PMN E. As shown in **Figure 7a**, all three intrabodies were expressed at high levels in the cytoplasm of transfected cells where they co-localized with cytoplasmic aggregates of TDP-43^ΔNLS^. There was no detectable reactivity with endogenous normal nuclear TDP-43, and there was no impact on the viability of the cells, suggesting that normal TDP-43 function was maintained.

**Figure 7.**
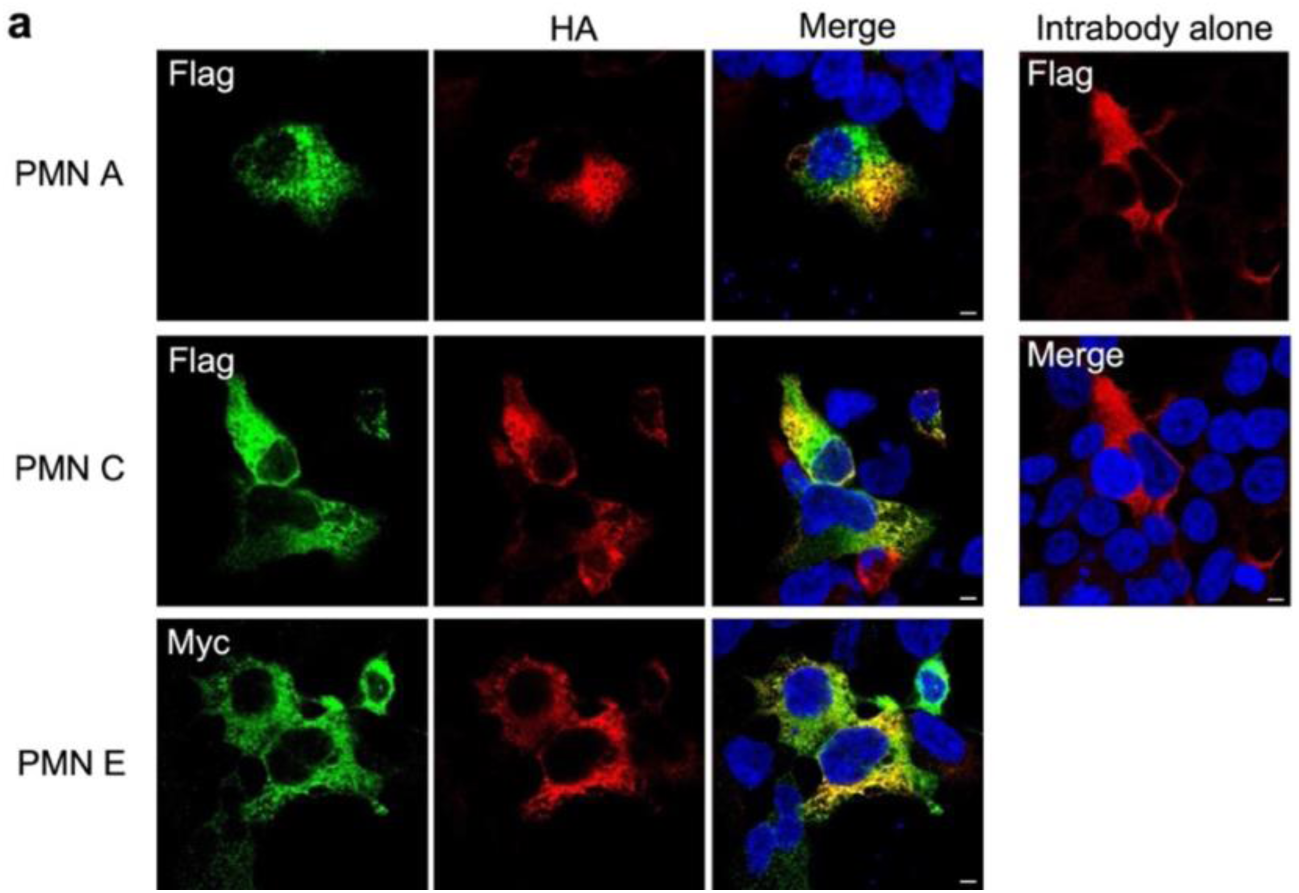
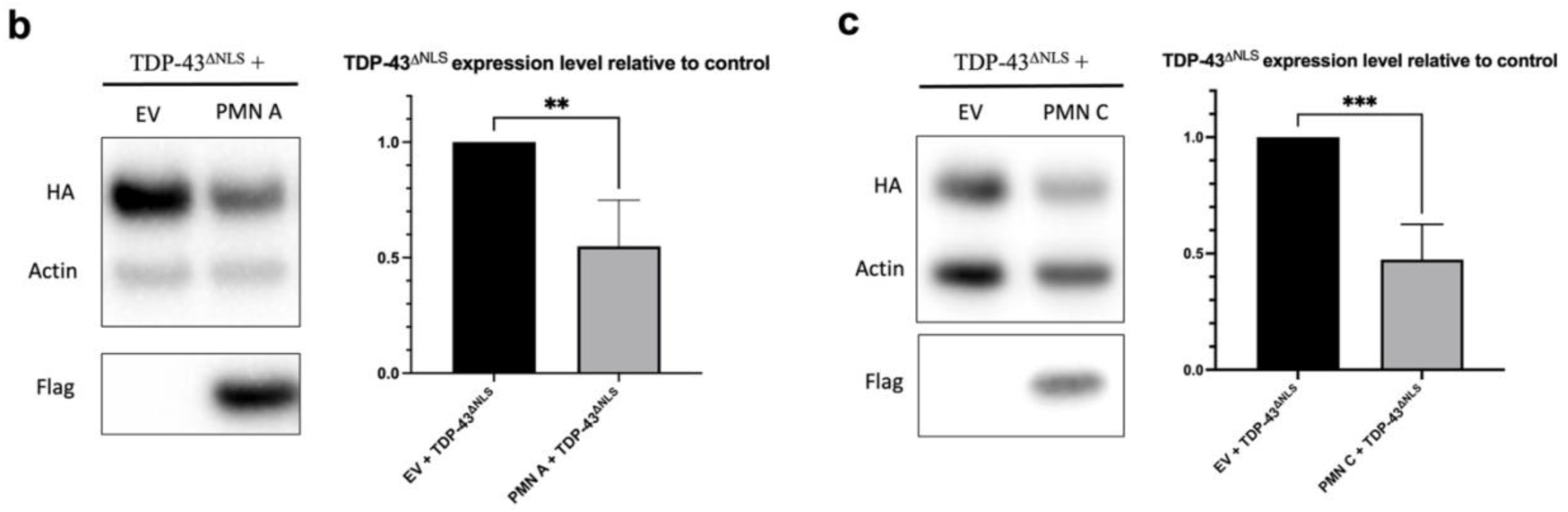
TDP-43 intrabodies bind to TDP-43^ΔNLS^ cytoplasmic aggregates and accelerate TDP-43^ΔNLS^ degradation. Immunocytochemical analysis demonstrates that transfected TDP-43 intrabodies (Flag, Myc-tagged) interact with cytoplasmic aggregates of TDP-43^ΔNLS^ (HA), and are exclusively localized to the cytoplasm when expressed alone (**a**). Scale bars: 10 μm. WB analysis shows PMN A and PMN C significantly accelerate TDP-43^ΔNLS^ degradation (**b, c**). *Statistics*, Student’s *t*-test, unpaired two-tailed. \*\**p ≤* 0.01, \*\*\**p ≤* 0.001. n=4

Second, to assess the ability of the intrabodies to clear misfolded aggregates of TDP-43^ΔNLS^, HEK293T cells were co-transfected with TDP-43^ΔNLS^ and PMN A, PMN C, or control empty vector (EV), followed by WB analysis. As shown in **Figures 7b & c**, PMN A and PMN C significantly reduced the level of intracellular TDP-43^ΔNLS^ by ∼50%, supporting the potential of intrabodies to promote the degradation of pathogenic TDP-43 aggregates within affected cells.

### TDP-43 intrabody alleviates the formation of SGs and TDP-43 aggregates upon chronic stress

As a more disease-relevant system, intrabody activity was also characterized in iPSC-MN lines from ALS patients carrying the TDP-43^N352S^ or FUS^R521G^ mutation. As shown above, the mAbs did not recognize TDP-43 in physiological SGs formed in response to acute oxidative stress (**Fig. 3**). SG proteins have been found to co-aggregate with TDP-43 in the spinal cords of ALS patients (34, 50–52), although pathological TDP-43 aggregates need not be formed in SGs (53). To better mimic the chronic stress conditions occurring during neurodegeneration, iPSC-MNs transduced with lentiviral vector expressing intrabody (PMN E), or luciferase as a negative control, were stressed with puromycin (5 μg/ml) for 24 hrs, followed by washout and recovery for 24 hrs. As shown in **Figure 8**, SGs were formed upon puromycin treatment, and dissolved during recovery in the ALS MN lines and a MN line derived from a healthy control (**Fig. 8a-c, h, j, & l**). Compared to the luciferase control, expression of the intrabody significantly reduced SG formation in the TDP-43^N352S^ and FUS^R521G^ ALS lines but not in the control MN line, presumably because of the absence of misfolded pathological TDP-43 in healthy neurons. The salutary effects of intrabody TDP-43 misfolding-specific PMN E in FUS^R521G^ suggests that NTD-unfolded wild-type TDP-43 may contribute to pathology in FUS-FALS. Interestingly, TDP-43 aggregates, or puncta, were detectable at baseline in the cytoplasm of TDP-43^N352S^ and FUS^R521G^ MN lines but not in control MNs. In ALS MN lines, the TDP-43 puncta were increased by puromycin stress and persisted with a further increase during recovery even when SGs had dissolved. The expression of PMN E reduced TDP-43 puncta at all stages (**Fig. 8e, f, k, & m)**, in agreement with its ability to target misfolded aggregates of TDP-43. TDP-43 puncta did not form in healthy control MNs, which are not expected to contain aggregation-prone TDP-43 (**Fig. 8d & i**). These results suggest that an intrabody selective for misfolded TDP-43 may have the potential to alleviate pathological SG formation and inhibit pathogenic TDP-43 cytoplasmic aggregation under the chronic stress conditions that occur in ALS.

**Figure 8.**
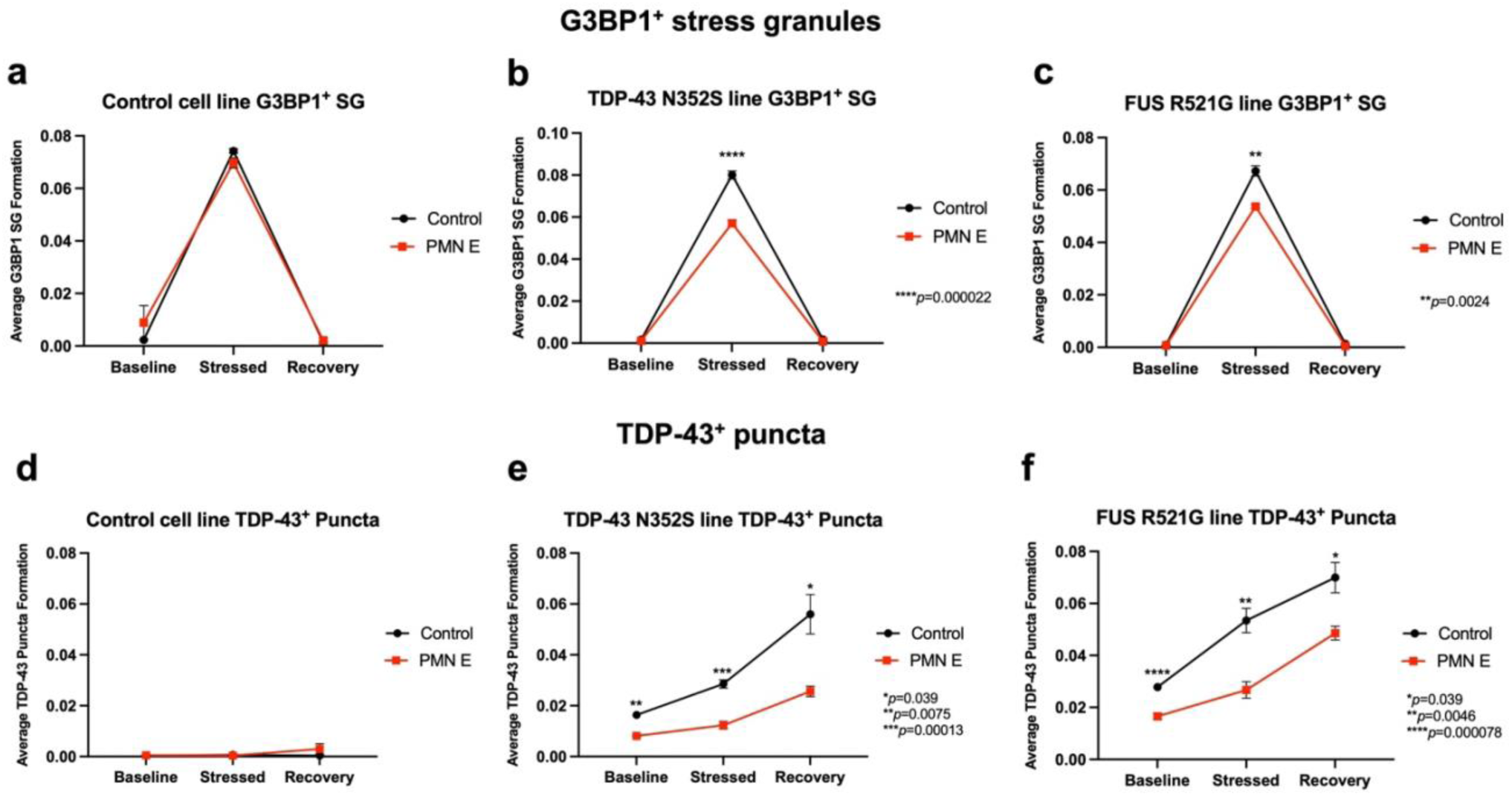
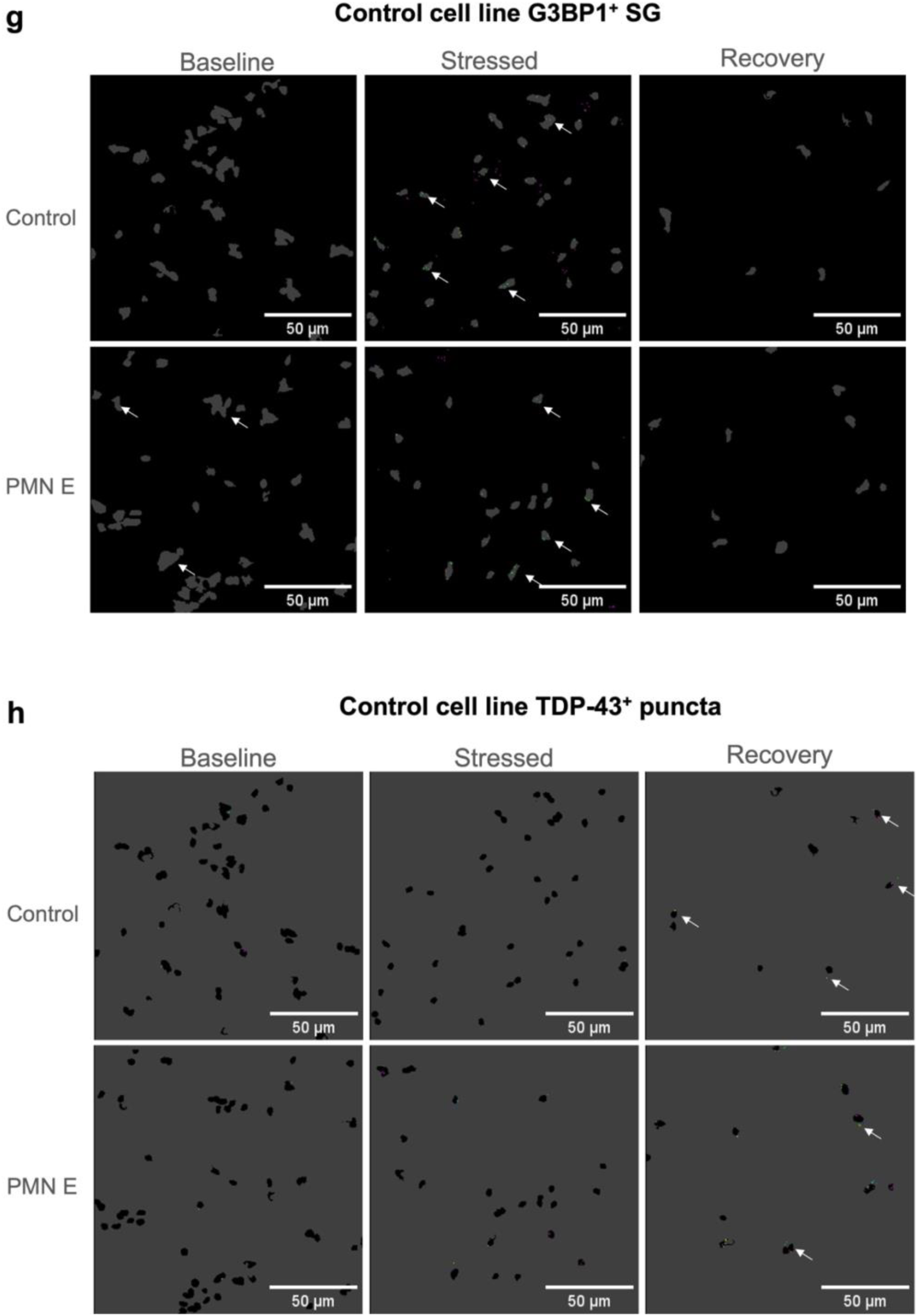
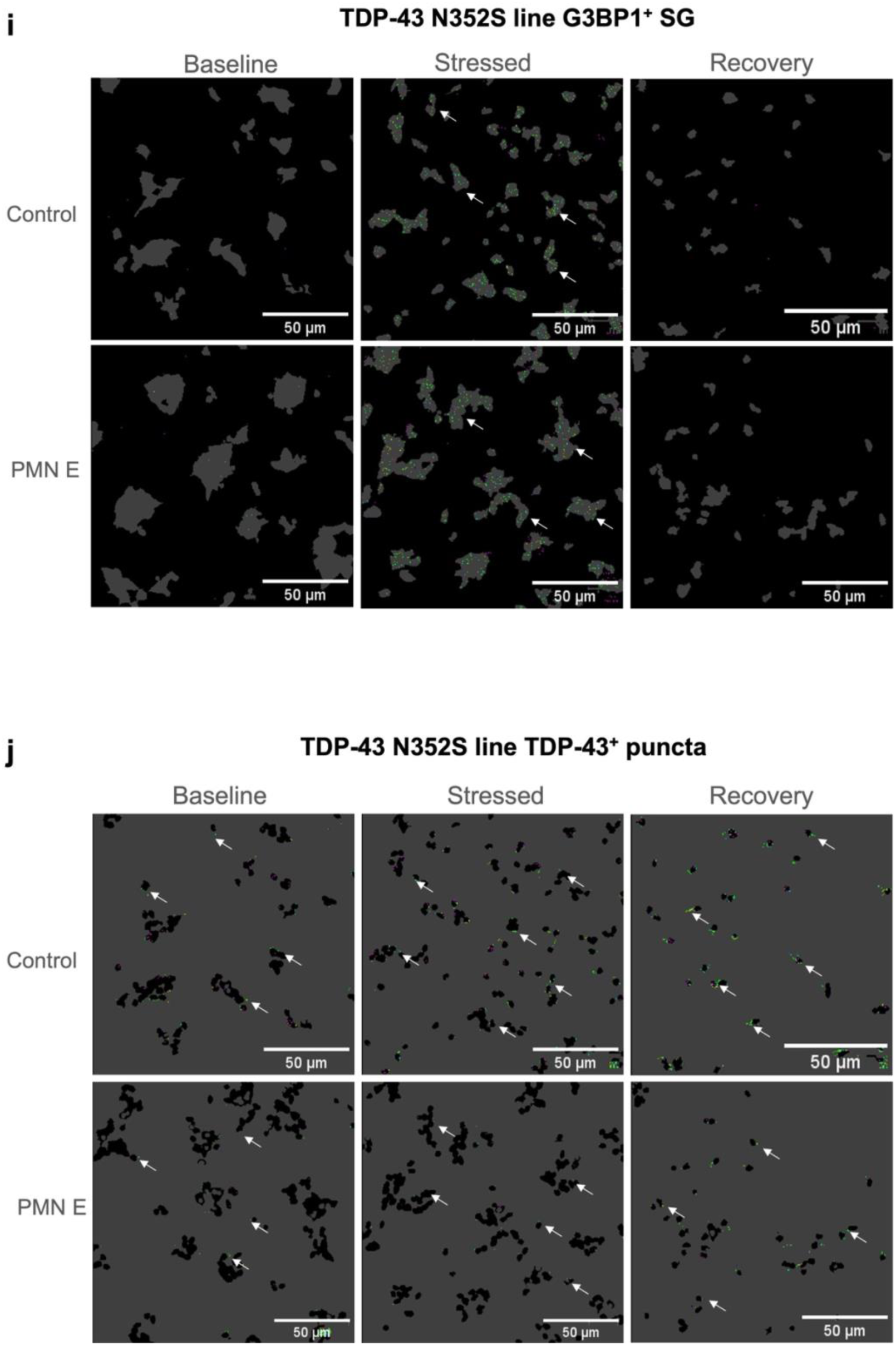
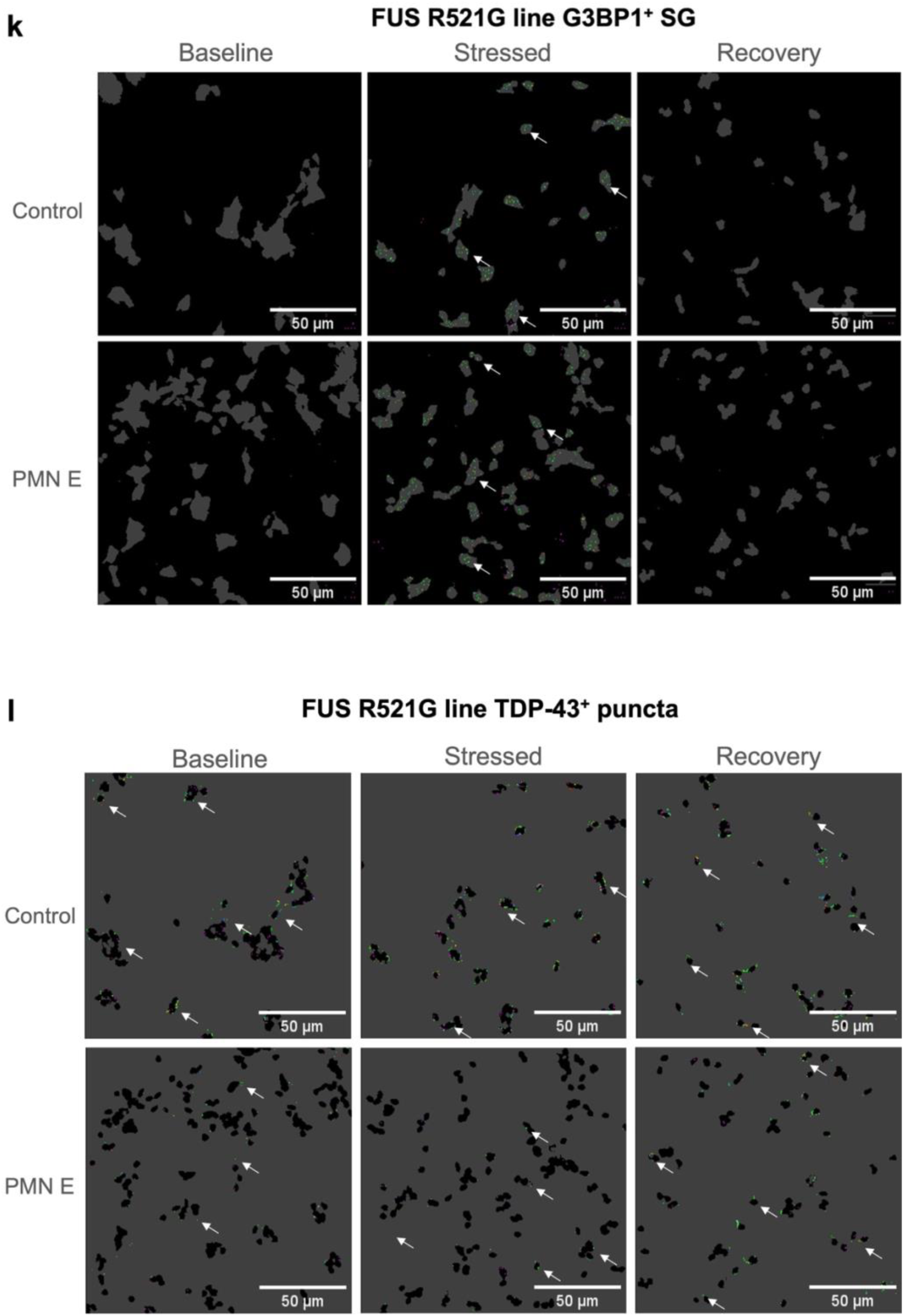
TDP-43 intrabody reduces SG formation and TDP-43 aggregates in iPSC-MNs under chronic stress conditions. Quantification of stress granules (G3BP1^+^ SG, a-c) and TDP-43 cytoplasmic aggregates (TDP-43^1^ puncta, d-f) upon puromycin stress (5 pg/ml, 24 hrs) and recovery following puromycin washout (24 hrs) in iPSC-MNs derived from healthy control, or patients carrying TDP-43^N352S^ or FUS^R^-^21G^ mutation transduced with lentiviral vectors expressing PMN E or luciferase as a control. Statistics, Multiple unpaired /-test. Error bars: SEM n=5-6. Representative CellProfiler images of control cell line (G3BP1^+^ puncta, g; TDP-43^+^puncta, h), TDP-43^N352S^ line (G3BPT puncta, i; TDP-43’ puncta, j), and FUS^R521G^ line (G3BP1* puncta, k; and TDP-43 puncta, 1). G3BP1 puncta are defined as the green dots sitting within the cell body mask, while TDP-43 puncta are defined as the green dots sitting outside of the nuclei mask. Pink dots denote speckles that were picked up by CellProfiler within the set intensity threshold but are not located in the correct area of the mask.

## Discussion

TDP-43 plays a major role in both neuronal physiological integrity and neurodegeneration. In ALS and FTLD-TDP, mutations in *TARDBP* are associated with a small percentage of disease, but wild-type TDP-43 cytoplasmic aggregation is observed in the vast majority of cases of ALS. With *TARDBP* mutations and wild-type TDP-43 being associated with ∼97% of all ALS, there is presumably an overlap of pathogenic pathways in genetic and non-genetic roles of TDP-43 in disease.

The pathogenic role of TDP-43 in ALS, FTLD-TDP and AD is believed to be due to a toxic synergy between loss of physiological functions (LoF) due to nuclear depletion, and gain-of-function (GoF) from cytoplasmic mislocalization and aggregation of TDP-43, with perhaps ∼50% relative contributions of LoF and GoF (54). LoF primarily results from defects in RNA splicing (55). GoF associated with cytoplasmic aggregates has been associated with neurodegeneration through a number of mechanisms, including mitochondrial and metabolic dysfunction (56), endoplasmic reticulum stress (57), defects in mRNA localization (58), and global protein translation (59, 60). Additionally, less well-recognized GoFs include: 1) prion-like recruitment of nascently translated TDP-43 to cytoplasmic aggregates (61); 2) transmission of prion-like seeds to adjacent cells and synaptically networked neurons (8); and 3) the co-aggregation of non-TDP43 proteins such as SOD1 that acquire prion-like misfolding and toxic propagation to other CNS cells, (62–64).

The NTD of TDP-43 is a well-conserved structural moiety that participates in LoF and GoF in disease (65). Deletion of the first 10 residues of TDP-43 negatively impacts the folding of the NTD to impair proper homodimerization and TDP-43-regulated splicing, and also renders the holoprotein more susceptible to cytoplasmic aggregation (66, 67). Our NTD epitope _65_DAGWGNL_71_, exposed by TDP-43 misfolding, was first identified by us to expose the Trp68 local sequence, which is buried and antibody-inaccessible in the normally folded NTD (22). Solvent-exposed Trp68 of misfolded TDP-43 was found to trigger SOD1 misfolding and prion-like propagation (22).

We believe that the exposure of our NTD epitope is also consistent with the observed increased proportion of dysfunctional monomeric TDP-43 in the nucleus. Trp68 is a key residue in the proper folding of the NTD, which consequently is responsible for NTD electrostatic oligomerization of TDP-43 for the nuclear splicing of pre-mRNA (65). Moreover, a peptide adjacent to our sequence in the NTD, _41_LRYRN_45_, is exposed in monomeric TDP-43, which has been demonstrated to lack pre-mRNA splicing activity (68, 69). Interestingly, it has been found that pre-mRNA splicing defects apparently precede neuronal dysfunction in ALS, in which neuronal cell death is more strongly associated with cytoplasmic aggregates (70).

Based on the current knowledge of TDP-43 biology and the findings described above, targeting of our misfolded TDP-43 NTD epitope has the potential to mitigate GoF defects arising from TDP-43 aggregation, as well as inhibit pathogenic recruitment of other proteins into cytoplasmic aggregates. In the present study, mAbs and intrabodies generated against this epitope recognized misfolded TDP-43 but not on properly folded normal TDP-43 (22). The mAbs showed high binding affinities to the epitope by SPR analysis and the reactivity and selectivity for pathogenic TDP-43 were verified in an *in vitro* cell system as well as in tissues from patients and normal controls. Functionally, the mAbs and intrabodies inhibited cell-to-cell propagation of misfolded TDP-43 and promoted degradation of intracellular aggregates in both a cell line and iPSC-MNs from ALS patients. These results provide proof-of-concept evidence supporting selective targeting of misfolded TDP-43 as a potentially safe and effective strategy for the treatment of neurodegenerative diseases associated with TDP-43 proteinopathy, such as ALS, FTD, and AD.

## Materials and Methods

### Generation of mAbs

Supernatants from mouse hybridoma clones and isolated rabbit B cell clones were tested for reactivity with the peptide by enzyme-linked immunosorbent assay (ELISA). Selected clones with high levels of binding were further characterized. The immunogen peptide was synthesized by CPC Scientific (Sunnyvale, CA, USA). mAbs were generated by ImmunoPrecise Antibodies (Victoria, BC, Canada).

### SPR

SPR kinetic analysis was conducted to assess the binding affinity of candidate mAbs to the TDP-43 N-terminal epitope peptide. BSA-conjugated peptide was immobilized on sensor chips at a low density of ∼50 response units (RUs). mAb1-mAb5 were diluted by 4-fold (31.25 nM, 7.81 nM, 1.95 nM, 0.98 nM, 0.24 nM) and injected over the surface. Binding curves were fitted to a Langmuir 1:1 interaction model.

### Cell culture, plasmids, and transfection

HEK293T cell line was purchased from American Type Culture Collection (ATCC, Rockville, MD, USA), and maintained in Dulbecco’s Modified Eagle Medium (DMEM) supplemented with 10% fetal bovine serum (FBS), GlutaMax^TM^ (2 mM) and antibiotics (50 U/ml penicillin and 50 mg/ml streptomycin) at 37°C in 5% CO_2_.

#### Plasmids

Human influenza hemagglutinin (HA)-tagged TDP-43^ΔNLS^ (K82A/R83A/K84A) was generated as previously described (63). PMN A scFv plasmid, which encodes the variable regions of the VH and VL chains of mAb1 joined by a linker and contains a Flag tag in the N-terminus and a lysosomal targeting signal in the C-terminus, was constructed by Creative Biolabs (Shirley, NY, USA). PMN C and E scFv plasmids, which encode the variable regions of the VH and VL chains of mAb5 joined by a linker, were constructed by Creative Biolabs and Syd Labs (Hopkinton, MA, USA), respectively. PMN C contains a Flag tag in the N-terminus and a lysosomal targeting signal in the C-terminus. PMN E contains a Myc tag in the C-terminus.

#### Transfection

cDNA plasmids were delivered into HEK293T cells using Lipofectamine LTX reagent (ThermoFisher Scientific, Waltham, MA, USA) following the manufacturer’s instruction. Cells were analyzed 48 hrs post-transfection.

### Immunocytochemistry

HEK293T cells were washed twice with phosphate-buffered saline (PBS) and fixed in 4% paraformaldehyde (PFA) for 15 min at room temperature (RT), followed by a wash with 20 mM glycine for 10 min at RT with constant rocking to quench residual PFA. Cells were then incubated with blocking buffer containing PBS, 1% Bovine Serum Albumin (BSA), 10% normal goat serum (NGS), and 0.1% Triton-X-100 for 30 min at RT. The following primary antibodies were incubated for 1 h at RT or overnight (ON) at 4°C: mouse monoclonal anti-misfolded TDP-43, mAb1 (10 μg/ml), rabbit monoclonal anti-misfolded TDP-43, mAb5 (2 μg/ml), rabbit polyclonal anti-HA (Abcam, Cambridge, UK, ab9110, 1:1,000),), rat monoclonal anti-HA (ThermoFisher, 11867423001, 1:1000), rabbit polyclonal anti-G3BP1 (Rosemont, IL, USA, 13057-2-AP, 1:1,000), rabbit monoclonal anti-Flag (Cell Signaling, Danvers, MA, USA, 2368S, 1:2000), rabbit polyclonal anti-c-Myc (Abcam, ab9106, 1:1,000). Cells were then washed with PBS/0.1% Triton-X-100 three times for 10 min with constant rocking, followed by incubation with Alexa Fluor^TM^ goat anti-rabbit 568, goat anti-mouse 488, or goat anti-rat 594 secondary antibody (ThermoFisher, 1:1,000) for 30 min at RT in the dark. Cells were then washed with PBS/0.1%Triton-X-100 three times for 10 min, dipped in 5% PBS, and mounted with ProLong Gold Anti-fade Reagent with DAPI (ThermoFisher, P36931). Cells were analyzed by confocal microscopy (Leica TCS SP8 MP, Wetzlar, Germany).

### Western blotting

Cells were washed twice with ice-cold PBS and lysed in 2% SDS, followed by sonication at 30% power for 15 sec to extract total protein. Protein content was determined by BCA assay (ThermoFisher). 10 µg of protein was separated on 4-12% NuPAGE Bis-Tris SDS-PAGE (ThermoFisher), transferred onto a PVDF membrane, and blocked in Tris buffered saline (TBS) containing 5% skim milk and 0.1% Tween-20 for 1 h at RT. The following primary antibodies were incubated ON at 4°C: rabbit anti-HA (Abcam, ab9110, 1:1,000), mouse anti-β-actin (Applied Biological Materials, Richmond, BC, Canada, G043, 1:1,000), mouse anti-GAPDH (ThermoFisher, AM4300, 1:100,000). Membranes were washed with TBS/0.1%Tween (TBST) three times for 10 min at RT with constant rocking, followed by horseradish peroxidase (HRP)-conjugated goat anti-mouse IgG (Sigma, St. Louis, MI, USA, AP181P, 1:5,000) or donkey anti-rabbit secondary IgG secondary antibody (Sigma, AP182P, 1:5,000) incubation for 30 min at RT. Membranes were then washed with TBST three times for 10 min, and developed with SuperSignal^TM^ West Femto Maximum Sensitivity Substrate (ThermoFisher, 34094).

### Immunohistochemistry

For mAb1 staining, brain samples were obtained from The Netherlands Brain Bank, Netherlands Institute for Neuroscience (Amsterdam, The Netherlands). Formalin-fixed paraffin-embedded tissue sections (8 μm) were mounted onto Superfrost plus tissue slides (Menzel-Gläser, Germany) and dried ON at 37°C. Sections were deparaffinized and subsequently immersed in 0.3% H_2_O_2_ in PBS for 30 min to quench endogenous peroxidase activity. The sections were then exposed to sub-boiling antigen retrieval buffer containing 10 mM Tris base, 1 mM EDTA solution, 0.05% Tween 20, pH 9.0. Primary antibodies mAb1 (1:2000) and rabbit polyclonal pTDP-43 (Cosmo Bio, Carlsbad, CA, USA, 1:8,000) were subsequently diluted in antibody diluent (Sigma, St. Louis, MO, USA) and incubated ON at 4°C. Secondary EnVision® horseradish peroxidase (HRP) goat anti-mouse antibody (ThermoFisher, 31430) was incubated for 30 min at RT, followed by 3,3-Diaminobenzidine (DAKO, Agilent Technologies, Santa Clara, CA, USA, S302283-2) incubation for 10 min. Sections were subsequently counterstained with hematoxylin to visualize the nuclei of the cells, dehydrated, and mounted using Quick-D mounting medium (Klinipath, Duiven, Netherlands).

For mAb5 staining, human spinal cord and brain samples were provided by Dr. Ian Mackenzie (University of British Columbia, Canada). Fresh-frozen sections (25 μm) were mounted onto charged glass slides, fixed by immersion in 10% neutral-buffered formalin (NBF) for 5 min, and washed extensively in Tris buffered saline (TBS, pH 7.4). Between subsequent steps, slides were washed in TBS containing 0.2% Triton-X-100. Endogenous peroxidase activity was quenched with 0.3% H_2_O_2_ in methanol. Sections were first blocked in TBS containing 10% NGS, 3% BSA, and 0.2% Triton-X-100, and further blocked with an avidin/biotin blocking kit (Vector Laboratories, Burlingame, CA, USA, SP-2001) following the manufacturer’s instructions. Primary antibodies rabbit monoclonal mAb5 (1 μg/ml) and rabbit polyclonal pan-TDP-43 (ProteinTech, 10782-2-AP, 1 μg/ml) were diluted in background-reducing antibody diluent (Agilent, S302283-2) and incubated ON at 4°C in a humidified chamber. Sections were subsequently incubated in biotin-conjugated F(ab)’2 fragment goat anti-rabbit IgG (H&L) secondary antibody (ThermoFisher, A-21246, 1:1,000) for 1 h, then labelled using the avidin/biotin complex-HRP system (Vector, PK-6100). Staining was visualized using a VIP HRP substrate kit (Vector, SK-4600). Sections were subsequently counterstained with methyl green (Vector, H-3402-500) to visualize the nuclei of the cells, dehydrated, cleared, and mounted using Permount (ThermoFisher, SP15-100).

### RT-QuIC assay

2 μl of 10 % brain homogenates diluted 10-fold were added to wells containing 98 μl of the master mix with final concentrations of 10 μg/ml recombinant TDP-43 (263-414), 20 mM urea, 20 mM Tris HCl pH 8.0, and 10 µM of ThT. TDP-43 (263-414) was filtered through a 100 kD MWCO filter (PALL #OD100C3, PALL Corporation, Puerto Rico) immediately prior to use. 10 μg/ml of TDP-43 antibodies were added before seeding with brain homogenates. Plates were sealed with clear adhesive sealing tape and placed in an Omega FLUOStar plate reader (BMG LABTECH, Ortenberg, Germany) pre-heated to 37 °C with no shaking. ThT fluorescence measurements (450 +/− 10 nm excitation and 480 +/− 10 nm emission; bottom read) were taken every 45 min. Experiments, each with triplicate wells, were repeated three times.

### Intercellular transmission

Intercellular transmission assay was performed following a previously established protocol (23). Briefly, donor cells were transfected with TDP-43^ΔNLS^ for 48 hrs. Conditioned medium was subsequently incubated with 30 μg/ml of mAb1 and control mIgG1 (BioLegend, San Diego, CA, USA, MG145), or mIgG2a (Abcam, ab185802) with constant rotation for 1.5 hrs at RT. The resulting medium was then added to naïve recipient cells and incubated for 48 hrs, followed by WB analysis to quantify the ability of mAb1 to inhibit TDP-43^ΔNLS^ intercellular transmission using Image Lab 5.2.1 to measure band density (Bio-Rad Laboratories, Hercules, CA, USA).

### Generation of motor neuron progenitors (MNPs)

iPSC-MNP generation was adapted from a previously published protocol (71). All iPSC lines were obtained and differentiated MNP stage, except *FUS R521G,* which was obtained and differentiated into MNPs from the iPSC stage. Briefly, MNPs were dissociated with Accutase (ThermoFisher, A6964) into single cells and plated at 1 x 10^6^ cells per well of a Matrigel (ThermoFisher, DLW356231)-coated 12-well plate. Matrigel-coated plates were made using 500 μl of Matrigel (1:60 v/v in DMEM) per well of a 12-well plate and incubated ON. Plated MNPs were grown for 24 hrs in DMEM/F12 supplemented with GlutaMax^TM^, B-27 (ThermoFisher, 17504044, 1:50) and N-2 (ThermoFisher, 17502048, 1:100). After plated, MNPs were grown for 24 hrs and transduced with lentivirus containing the intrabody sequences at 10 μl per well as a pre-treatment setup.

### Generation of Lentivirus

To transduce the misfolded TDP-43-targeting intrabody into iPSC-MNs, the open-reading frame of the intrabody was cloned into individual lentiviral vectors for potential lentiviral transfection and transduction. After amplifying the intrabody sequence *via* polymer chain reaction, the amplicon was cloned into a pXR002: EF1a-dCasRx-2A-EGFP [gift from Dr. Patrick Hsu, University of California, Berkeley; Addgene (Watertown, MA, USA, 109050)] plasmid backbone utilizing restriction sites, *EcoRI* and *PacI*. With the amplicon situated downstream of an Ef1α promoter sequence and attached to a gBlock^TM^ (IDT, Coralville, IA, USA) consisting of an IRES-mCherry-V5 sequence, the final plasmid was generated using Gibson assembly and sustained in Stbl3 *E. coli* competent cells (ThermoFisher, C737303). Sanger sequencing was used to validate the individual clones, and the DNA was prepared *via* the ZymoPURE II Plasmid Maxiprep kit (Zymo Research Corp, Irvine, CA, USA, D4204) with endotoxin removal for eventual lentiviral packaging. Utilized primers for creating the constructs can be found in **Table 2**.

**Table 2.**
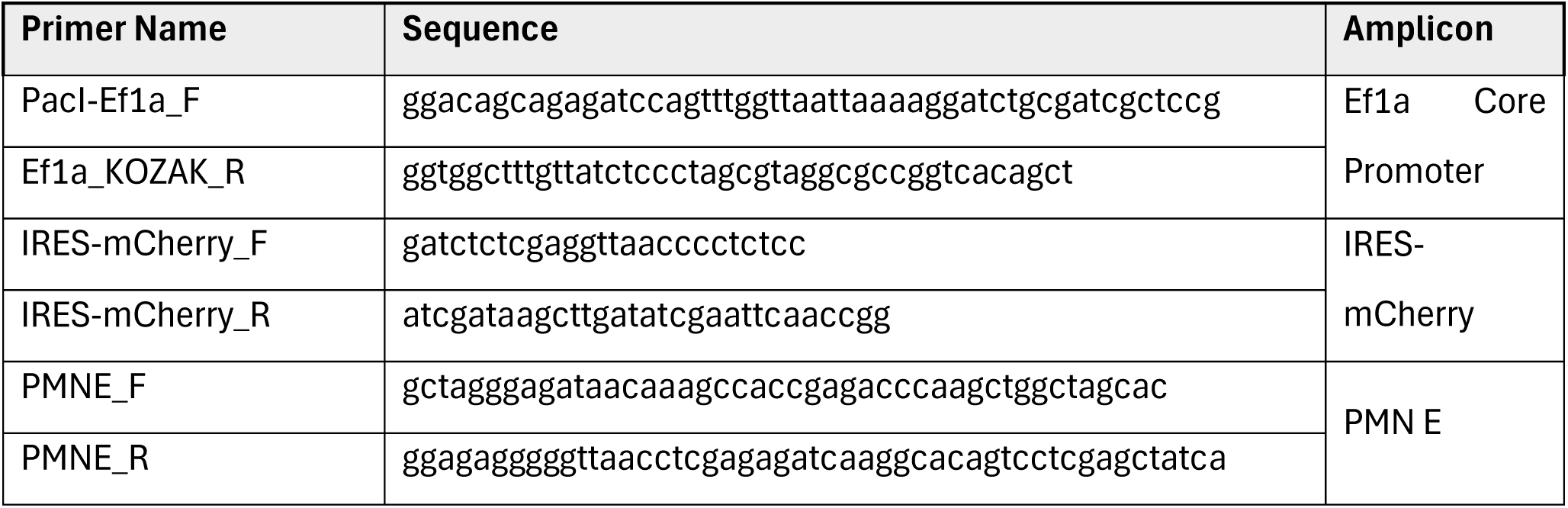

24 hrs prior to transfection of the constructed plasmid, HEK293T cells were seeded onto pre-coated 15 cm^2^ tissue culture dishes with 0.005% Poly-D-lysine (PDL) at 30-40% confluency and incubated at 37°C and 5% CO_2_ in DMEM containing 10% FBS without antibiotics. Once the cells reached 70-90% confluency the following day, 20 µg lentiviral plasmid, 6 µg pMD2.G (VSV-G) envelope expressing plasmid (gift from Dr. Didier Trono, University of Geneva, Switzerland; Addgene, 12259), and 12 µg psPAX2 packaging plasmid (gift from Dr. Didier Trono; Addgene, 12260) were added to the cells for transfection using Lipofectamine 3000 (ThermoFisher, L3000001) following the manufacturer’s instructions. 6 hrs following transfection, a full media change was performed and replaced with DMEM containing 20% FBS without antibiotics. The lentivirus-containing media was collected at 48- and 72 hrs after transfection and stored at 4°C. Lenti-X Concentrator reagent (Takara Bio, Kusatsu, Japan, 631231) was used following the manufacturer’s instructions to treat the lentiviral supernatants. Following treatment, pellets were resuspended using 1/100 of the original collection volume in non-supplemented DMEM. Lentivirus was then aliquoted and stored at -80°C until utilized for transduction.

### Plate Coatings for Cell Culture Maintenance and Screening

For culturing iPSCs and MNPs, 10 cm^2^ tissue culture dishes were coated with 5 ml of Matrigel (1:60 v/v in DMEM) and incubated at 37°C for 60 min. For differentiating motor neurons, 6-well and 384-well plates were coated with 1 ml and 20 μl, respectively, of a combined aqueous mixture of 0.001% (w/v) PDL hydrobromide and 0.001% (w/v) Poly-L-ornithine (PLO) hydrobromide and incubated at 37°C ON. The PDL/PLO solution was then aspirated from the plates, followed by washing twice using Dulbecco’s phosphate-buffered saline (DPBS), coated in 1 ml and 20 μl of mouse laminin (MilliporeSigma, L2020-1MG, 1:50 v/v in DPBS) for 6-well plates and 384-well plates, respectively, and incubated at 37°C ON. Before seeding, laminin was aspirated from the plates.

### Puromycin Stress Assay

Matured motor neurons were seeded onto PDL/PLO and laminin (1:50 v/v) coated Perkin-Elmer 384-well plates at 2 x 10^4^ cells per well. A complete media change was performed the following day. On the second day after seeding, the cells were stressed with puromycin (5 μg/ml) for 24 hrs. After 24 hrs, cells were either fixed in 4% PFA at RT for 1 h to keep at stressed condition or three half washes were performed, with 5 min incubations at 37°C after each wash to stimulate recovery. After a 24-hr recovery, cells were fixed in 4% PFA at RT for 1 hr. Following fixation, four times of 5 min half-washes with DPBS were performed at RT. Cells were then permeabilized and blocked in DPBS containing 0.1% Triton-X-100 and 5% NGS. Primary antibodies mouse monoclonal anti-TDP-43 (Abnova, Taipei, Taiwan, H00023435-M01, 1:1000) and rabbit polyclonal anti-G3BP1 (MBL, RN048PW, 1:500) were incubated at 4°C ON, followed by eight times of 5 min half-washes in PBS/0.01% Triton-X-100 at RT on a platform rocker. Secondary antibodies Alexa Fluor^TM^ goat anti-rabbit 647 and goat anti-mouse 488 (ThermoFisher, 1:500) were then applied for 2 hrs at RT in the dark, followed by eight times of 5 min half-washes in PBS/0.01% Triton-X-100 at RT, all on a platform rocker. DAPI (Thermo Fisher, 622481:10,000 v/v) was applied for 10 minutes at RT in the dark, followed by a 10 min half-wash in PBS/0.01% Triton-X-100 at RT on a platform rocker. Finally, cells were mounted with glycerol (MilliporeSigma, G9012-2L, 50% v/v) in PBS.

#### Robotic Imaging

384-well plates were imaged *via* a Nikon widefield microscope system (Nikon Eclipse Ti2-E) using an NIS-Elements software (version 5.42.03 High content analysis package). Eight fields of view, four at each corner of the well and four at each well’s center, were imaged with a 40x (NA 0.95 air) and across 7 z-stacks of 0.6µm thickness across a 3.6µm range for DAPI, G3BP1-Cy5, and TDP-43-G3BP1 at 460 nm, 635 nm, and 535 nm, respectively.

#### Automated Segmentation and Quantification of Images

384-well plate images were segmented and quantified using a custom CellProfiler pipeline as previously reported with modifications (71). Nuclei were identified and segmented using the DAPI images for the diameter. Cell bodies were then determined by overlaying the G3BP1-Cy5 images and drawing from the nuclei outwards to the edges of the G3BP1-Cy5 signal. To eliminate background imaging artifacts, such as dead cells or background fluorescence, the cell bodies were utilized as masks. Following masking, an image processing enhancement feature for speckle-like features was utilized to highlight punctate structures. These structures proceeded to be labelled as either G3BP1-Cy5 SGs or TDP-43-GFP SGs. The total image area was calculated for each of the identified features (overlap of features, punctate structures, and nuclei) and output as spreadsheets.

#### Statistical Analysis of Screen Assay Data

Intrabodies were deemed as effective in reducing SG formation by decreasing the ratios of the average area occupied by G3BP1 SGs divided by the average area occupied by the nuclei and the average area occupied by TDP-43 SGs divided by the average area occupied by the nuclei when comparing with the luciferase control in the stress condition, and if applicable, in the recovery condition, as well.

### Statistics

Statistical analysis was performed using GraphPad Prism 9 or 10 (GraphPad Software, San Diego, CA, USA).

## Declarations

### Ethics approval and consent to participate

All of the human post-mortem materials used in this study were obtained from subjects who had been evaluated at the Netherlands Brain Bank and the University of British Columbia Clinic for Alzheimer’s Disease and Related Disorders. All donors have given informed consent for autopsy and the use of the material and clinical information for research purposes in accordance with respective ethical review boards.

### Availability of data and materials

The datasets generated in the current study are available from the corresponding author upon reasonable request.

### Authors’ contributions

B.Z., B.W.C, G.W.Y., J.M.K., and N.R.C. designed the experiments. B.Z., J.A.C., S.L., E.G., M.Y.H., R.J.M., S.A., P.A., M.L.D and A.A.D. performed the experiments and analyzed the results. The manuscript was drafted by B.Z. and edited by M.Y.H., R.J.M., S.A., I.R.M., P.A., B.W.C., G.W.Y., J.M.K., and N.R.C. All authors reviewed the manuscript.

